# Scale-Dependent Coding of the Hippocampus in Relational Memory

**DOI:** 10.1101/2024.01.29.577883

**Authors:** Wei Ding, Yijing Lin, Bo Zhang, Jia Liu

## Abstract

Memory, woven into the very fabric of consciousness, serves as a time-traveling vessel in the mind, using detailed recollections not just for nostalgic reflection but as an abstract map for charting unknown futures. Here, we employed fMRI to investigate how the hippocampus (HPC) encodes detailed experiences into abstract knowledge (i.e., formation) and uses this knowledge for decision making (i.e., utilization) when human participants learned and then utilized spatial-temporal relations in a relational memory task. We found a functional gradient along the anterior-posterior axis of the HPC, characterized by representational similarity and functional connectivity with the autobiographical network. Here, the posterior HPC was more actively engaged in memory formation, whereas the anterior HPC was predominantly involved in memory utilization. Our computational modeling of relational memory further established a causal link between this functional gradient and the HPC’s well-documented anatomical gradient, as optimal task performance arose from a combination of a fine-grained representation of past experiences by the posterior HPC and a coarser representation of abstract knowledge for future planning by the anterior HPC. This scale-dependent coding scheme led to the emergence of grid-like, heading direction-like, and place-like units in our neural network model, analogous to those discovered in biological brains. Taken together, our study revealed the HPC’s functional gradient in representing relational memory, and further connected it to the anatomical gradient of place cells, supporting a unified framework where both spatial and episodic memory rely on relational representations that integrate spatial localization with temporal continuity.

## Introduction

The core function of episodic memory is to remember the past and predict the future (Schacter & Addis, 2007; Addis et al., 2007; Addis et al., 2009; Schacter et al., 2012; Schacter & Madore, 2016; Gershman, 2017; Josselyn & Tonegawa, 2020; Namboodiri & Stuber, 2021). This process apparently relies on different levels of precision during encoding and retrieval stages. Encoding is often tasked with meticulously recording intricate details, necessitating fine-grained representational capacity to capture the nuanced differentiation of individual experiences (Wixted et al., 2018; Borders et al., 2022). In contrast, the retrieval process, especially for decision-making, tends to utilize a more generalized and abstracted memory schema, as this coarse form of recall provides the essential framework required to navigate through decision-making processes without the encumbrance of excessive detail (Evensmoen et al., 2015; Audrain & McAndrews, 2022). However, it is unclear how the brain accomplishes this fascinating interplay between the granular encoding and the abstracted retrieval that underscores the versatility and sophistication of our memory systems.

Recently, a unified theoretical framework has been proposed to associate episodic memory with spatial memory in the hippocampus (HPC) (Cohen & Eichenbaum, 1993; Eichenbaum, 2000; Eichenbaum & Cohen, 2014), evidenced by the abstract cognitive map (Park et al., 2020), cognitive graph (Peer et al., 2021), schema (Farzanfar et al., 2023), and geometry (Nieh et al., 2021). This framework offers a novel perspective on the HPCs role in memory encoding and retrieval by integrating episodic and spatial memory into a cohesive concept of relational memory. Interestingly, a representational gradient of the HPC in spatial scale along its anterior-posterior axis has been identified (Kjelstrup et al., 2008; Brun et al., 2008; Peer et al., 2019). Specifically, place cells in the anterior HPC of primates (or ventral HPC of rodents) show larger receptive fields than those in the posterior HPC (or dorsal HPC of rodents) (Poppenk et al., 2013; Collin et al., 2015; Robinson et al., 2015; Brunec et al., 2018). In addition, functional divisions of labor along this representational gradient have also been identified. The anterior HPC is more engaged during autobiographical memory retrieval (Cabeza & St Jacques, 2007; McCormick et al., 2015; Mazzoni et al., 2019). In contrast, the posterior HPC is preferentially implicated in both short-term (Fernandez et al., 1998; Giovanello et al., 2009; Woolley et al., 2015; Zeithamova & Bowman, 2020) and long-term learning (Maguire et al., 2000; Weisberg et al., 2019; Weisberg & Ekstrom, 2021). Considering these anatomical and functional divisions, and the different precision levels required between encoding and retrieval stages, we hypothesize that the posterior HPC, with its smaller receptive fields and therefore higher precision, may be the site for fine-grained encoding of detailed experiences into abstract representations (i.e., abstract knowledge). In contrast, retrieval of this abstract knowledge may occur in the anterior HPC, where the larger receptive fields imply a lower precision.

To test this hypothesis, we utilized a relational memory (RM) task (Insa-Cabrera et al., 2011; Tartaglia et al., 2017), where human participants were tasked with mentally constructing and utilizing a sequence of spatial positions to predict subsequent positions (Fig. 1). In the task, participants learned the temporal structure of spatial positions through feedback on random guesses in the beginning steps of each trial (i.e., the relation formation state). Subsequently, they applied this learned sequence to accurately predict each position in the sequence in the later steps of each trial (i.e., the utilization state). Therefore, this task design allowed separate analysis of neural representations during memory formation and utilization states. In line with our hypothesis, we found greater involvement of the posterior HPC in the formation state, while the anterior HPC was more engaged in the utilization state, aligning with the representational gradient along the anterior-posterior axis of the HPC. To delve deeper into the underlying mechanism, we developed a hybrid neural network model combining convolutional and Long Short-Term Memory (LSTM) components to examine the impact of spatial scale on the formation and utilization states of memory. Computational modeling established a causal link that the relational memory’s effectiveness hinged on fine-grained encoding and coarse retrieval representations. In short, our study suggests that the HPC exhibits multi-scale representations along the anterior-posterior axis tailored to computational demands of memory at different stages.

**Figure 1.**
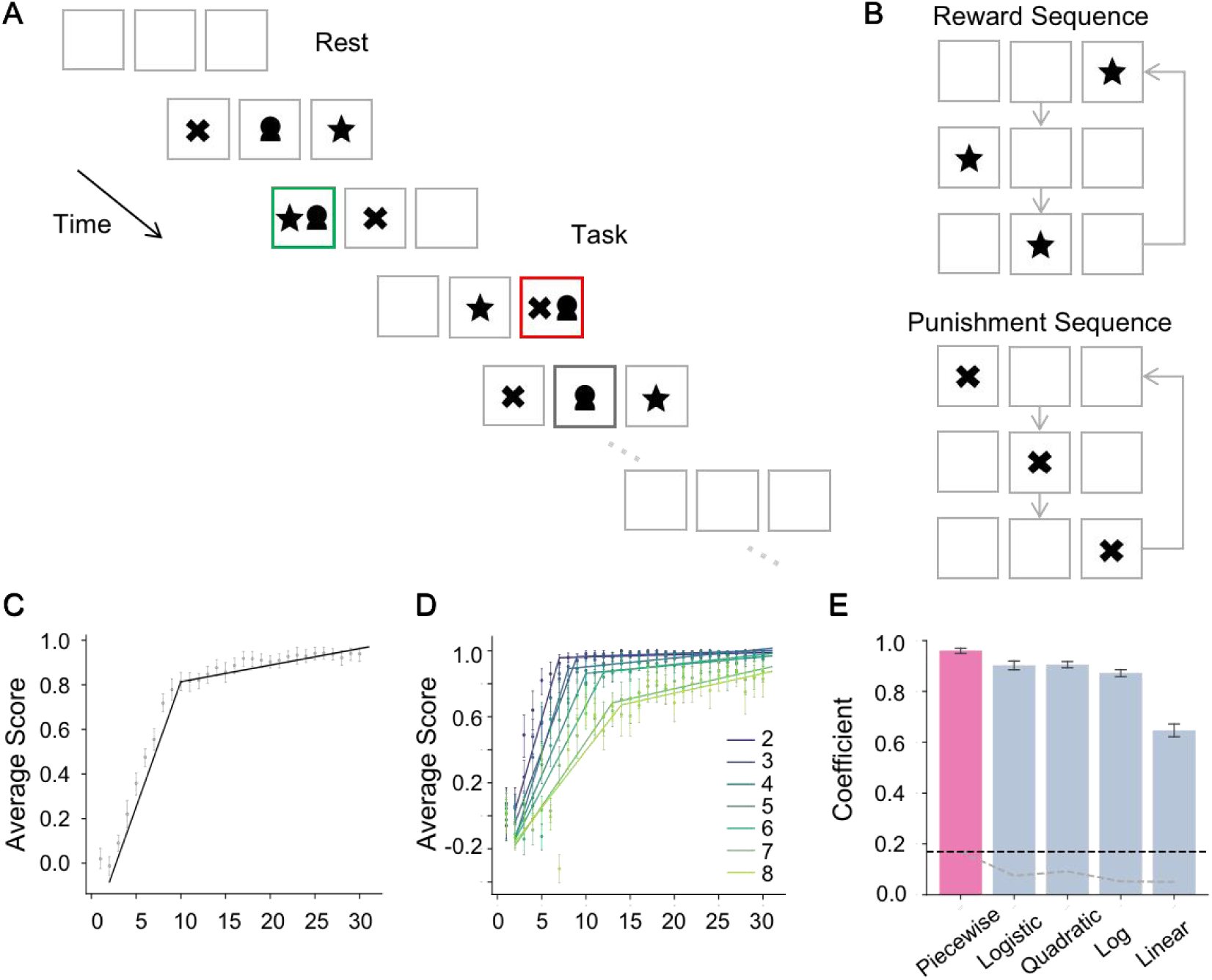
Experimental paradigm and behavioral performance. (A) Experimental paradigm of the RM task. A reward was presented when correctly placing the *Self* marker (avatar) in the same box (green) with the next *Good* cue (star), while a punishment was given when incorrectly placing the *Self* marker in the same box (red) with the next *Bad* cue (cross). Participants were encouraged to maximize reward and avoid punishment. (B) Exemplar sequences for the *Good* and *Bad* cues. The length of numeric strings is 3. The *Good* cue shows a [Right, Left, Middle] spatiotemporal sequence, while the *Bad* cue shows a [Left, Middle, Right] sequence. (C) Behavioral performance. Behavioral performance, averaged across sequences and participants, was improved as a function of increased steps. (D) The piecewise-like pattern, indexed by solid lines, was observed in all tested lengths of numeric strings. (E) The highest fitness to the behavioral performance was the piecewise linear function, as compared to the logistic, quadratic, logarithmic and linear functions. The dashed grey line denotes the uncorrected threshold of permutation, while the dashed black line denotes the corrected threshold with multiple comparisons. Error bars represent standard errors across participants.

## Results

### Memory formation and utilization states in relational memory task

A series of numeric strings, each composed of digits drawn from the set [1, 2, 3] and varying in length from 2 to 8, was constructed to represent sequences of locations for *Good* cues (denoted by a star) and *Bad* cues (denoted by a cross) within a single iteration, which were allocated to three horizontally aligned boxes on the screen (Fig. 1A). In a trial, sequences for both *Good* and *Bad* cues, derived from different numeric strings but of equal length, were iteratively presented until reaching a total length of 30 steps, respectively (Fig. 1B). Participants were instructed to place a *Self* marker (denoted by an avatar) at the anticipated location of the next *Good* cue while avoiding the location of the *Bad* cue, utilizing feedback from preceding steps. Participants’ performance was assessed by calculating the number of correctly placing the *Self* marker in the box containing the *Good* cue (i.e., reward) minus the number of incorrectly placing in the box with the *Bad* cue (i.e., punishment). Through this process, participants first learned the spatiotemporal relations of both *Good* and *Bad* cues, then applied this knowledge to predict the locations of the next *Good* and *Bad* cues. Participants’ behavioral performance exhibited a characteristic piecewise-like progression, initially manifesting a rapid rise in accuracy from random guessing, followed by a plateau of correct responses (Fig. 1C). This behavioral pattern was observed in all tested lengths of numeric strings, though a noticeable decrease in accuracy and a delay in the transition point were observed with increasing length of the numeric strings (Fig. 1D). To quantify the observation, we evaluated the fitness of a piecewise linear function to participants’ behavioral performance, against various functions such as logistic, quadratic, logarithmic, and linear functions, using the coefficient of determination ( *R^2^*) as the metric. Consistent with the observation, the piecewise linear function showed the highest fitness to the behavioral performance (Fig. 1E; *M*= 0.96, *SD* = 0.26; *p*< 0.05, corrected for multiple comparisons using permutation) compared to the alternative functions (logistic: [*M*= 0.90, *SD*= 0.04]; quadratic: [*M*= 0.90, *SD*= 0.03]; logarithmic function [*M*= 0.87, *SD*= 0.04]; and linear function: [*M*= 0.64, *SD*= 0.06]). This piecewise pattern suggests the existence of two distinct memory states with the task: one primarily pertaining to memory formation and the other to its subsequent utilization.

### Neural correlates of memory formation and utilization

Upon identifying two distinct memory states in the RM task, we further replicated previous research (Sullivan Giovanello et al., 2004; Olson et al., 2006; Hannula et al., 2006; Giovanello et al., 2009; Schwarb et al., 2016; Zheng et al., 2024) demonstrating hippocampal activation in relational integration using representational similarity analysis (RSA; Kriegeskorte et al., 2006) and functional connectivity (FC) approaches. In the RSA analysis, we calculated the multivoxel patterns of HPC activity during the RM task for each trial and constructed a correlation coefficient matrix across trials for each voxel (see Methods). We found that all subregions of the HPC, including Dentate Gyrus, Cornu Ammonis, and Subiculum, showed significantly higher representational similarity in comparisons to the resting state (Fig. S1; initial threshold: *p* < 0.01, corrected for multiple comparisons by permutation). In the FC analysis, we designated the HPC as a seed region and computed Pearson’s correlation coefficients between its averaged time course and that of other brain regions during the RM task (see Methods). We found significantly enhanced FC between the HPC and the autobiographical network (Svoboda et al., 2006; Burianova et al., 2010), including the Medial Prefrontal Cortex, Posterior Cingulate Cortex, and Angular Gyrus, during the task state in comparison to the resting state (Fig. S2; initial threshold: *p* < 0.01, corrected for multiple comparisons by permutation). Taken together, these findings align with previous research (Giovanello et al., 2009; Huijbers et al., 2011; Raichle, 2015; Smallwood et al., 2021), confirming the role of the HPC in relational memory processing during the RM task.

To investigate the neural representations of the HPC associated with the formation and utilization of relational memory, the BOLD signal series of the RM task was segregated into the formation and utilization states, which were approximately demarcated to have equal temporal durations (see Methods). An RSA analysis was performed to extract the neural representation profiles for the resting, formation, and utilization states, respectively (Fig. 2A). As expected, the representational similarity in the HPC was significantly higher in both the formation and utilization states than that in the resting state (Fig. 2B & E, initial threshold of *p*= 0.01, corrected for multiple comparisons by permutation).

**Figure 2.**
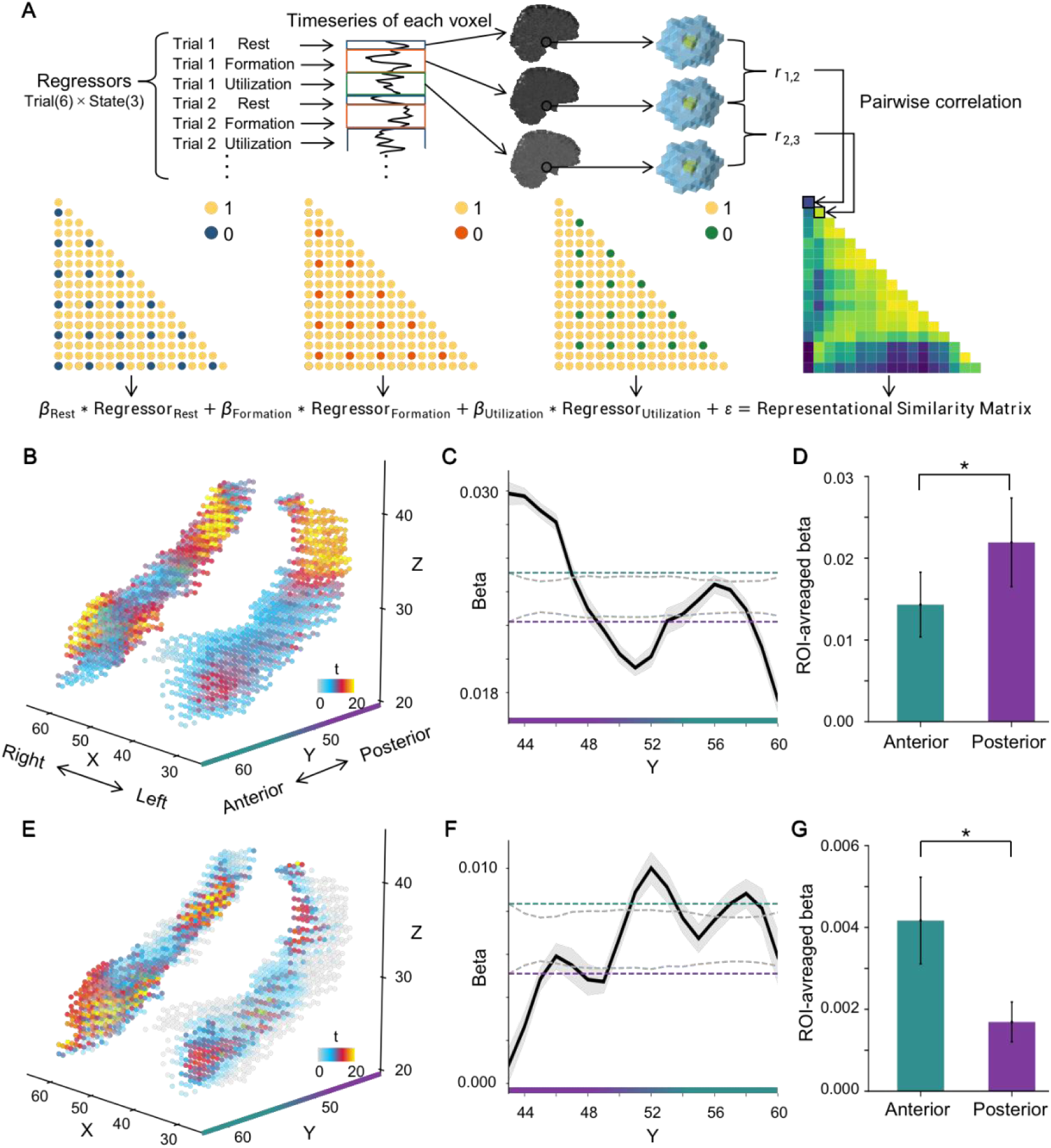
Function gradient in the HPC. (A) Schematic overview of RSA analysis. Multi-voxel patterns of the resting, formation and utilization states were extracted by a general linear model (GLM). The similarity matrix was constructed using a searchlight-based Pearson’s correlation of the multi-voxel patterns among trials. This matrix was fitted with another GLM with states specified by binary regressors. Within the binary matrix, blue, brown, and green points denote the “same” state. The estimated parameters of the GLM were then sent to the second-level analyses. (B) Neural representation of formation state in the HPC. Each point represents a voxel in the left and right hippocampus, respectively. The value at each point, indexed by t-value, reflects the contrast in representational similarity from the memory formation relative to the resting state. The X-, Y-, and Z-axes denote the MNI coordinates of the HPC. Specifically, the Y-axis corresponds to the hippocampal long axis, with green and purple colors representing the anterior and posterior HPC, respectively. (C) Variation in representational similarity along the hippocampal long axis during memory formation. The black line represents the averaged similarity across participants, with the shaded area denoting standard error. Gray dotted lines indicate the uncorrected thresholds derived from the 5^th^ and 95^th^ percentiles of the permutation distribution at each coordinate of the Y-axis, whereas green and purple dotted lines indicate the corrected threshold defined by the maximum of uncorrected thresholds. (D) ROI analysis of representational similarity between the anterior and posterior HPC. The representational similarity of memory formation state was significantly higher in the posterior HPC than that in the anterior HPC. (E – G) Neural representation of memory utilization state in the HPC. *: *p*< 0.05; Error bars: standard error.

Interestingly, the peak of the representational similarity in the formation state was located at the posterior HPC (MNI coordinates = [57, 45, 36], *t*(15) = 6.11, *p*< 0.001, two-tailed), while in the utilization state, it was identified in the anterior HPC (MNI coordinates = [58, 57, 26]; *t*(15) = 4.05, *p* < 0.001, two-tailed). This suggests an association between the functional dissociation in memory states and the anatomical gradient in the HPC. To quantify this observation, we averaged the representational similarity along the y axis of MNI coordinate system, which produced a vector demonstrating the variation in representational similarity along the hippocampal long axis for each memory state. First, a reverse pattern was observed between the two memory states, with higher representational similarity of memory formation in the posterior HPC and for memory utilization in the anterior HPC (Fig. 2C & F; initial threshold *p*= 0.01, corrected for multiple comparisons by permutation). Second, we partitioned the HPC into three equal volumetric portions along its long axis (Tompary & Davachi, 2017) and then performed an ROI analysis to explore the relationship between memory state and hippocampal gradient. We found that the posterior HPC showed significantly higher representational similarity as compared to the anterior HPC during memory formation (Fig. 2D; *t*(15) = 2.59, *p* < 0.05, two-tailed). Conversely, the anterior HPC showed significantly higher representational similarity as compared to the posterior HPC during memory utilization (Fig. 2G;*t*(15) = 2.66, *p* < 0.05, two-tailed). These findings illustrated a functional division of labor along hippocampal gradient for the two memory states in relational memory, suggesting that memory formation particularly relies on the fine-grained representation in the posterior HPC, while memory utilization preferentially involves the coarser representation in the anterior HPC.

Given the aforementioned evidence of the HPC’s division in representational similarity and its FC with the autobiographical network during the MR task, we predicted that the posterior HPC should predominantly connect with the autobiographical network during the formation state, while the anterior HPC should show stronger FC with this network during the utilization state.

To test this conjecture, we analyzed the FC between the HPC and the autobiographical network during the formation and utilization states of the RM task, respectively. Note that the medial temporal lobe was excluded from this analysis to avoid overlap with the HPC. We found that the posterior HPC showed stronger FC with the autobiographical network during the formation state (Fig. 3A; initial threshold *p*= 0.01, corrected for multiple comparisons by permutation), with the FC peak located at the coordinate [55, 52, 27] (*t*(15) = 4.98, *p* < 0.001, two-tailed). Conversely, during the utilization state, stronger FC with the autobiographical network was found in the anterior HPC (Fig. 3B; initial threshold *p*= 0.01, corrected for multiple comparisons by permutation), with the peak at [57, 61, 23] (*t*(15) = 3.29, *p*< 0.001, two-tailed). To quantify this observation, a contrast analysis between the utilization and formation states was performed on FC along the hippocampal long axis. We found that the FC difference between the two states shifted from negative to positive along the axis, moving from the posterior to anterior HPC (Fig. 3C; initial threshold *p* = 0.01, corrected for multiple comparisons by permutation). This connectivity gradient along the hippocampal long axis was further confirmed by a significant two-way interaction between the memory state (formation versus utilization) and the hippocampal gradient (posterior versus anterior) by an ROI analysis (Fig. 3D; *F*(1, 15) = 9.03, *p* = 0.004). *Post-hoc* t-tests showed that in the anterior HPC, the FC during the utilization state was significantly higher than that of the formation state (*t*(15) = 5.44, *p* < 0.001, two-tailed), while a reverse pattern, though not significant, was found in the posterior HPC (*t*(15) = 1.55, *p* = 0.14, two-tailed). In summary, the FC analysis also reveals a connectivity gradient of the HPC with the autobiographical network during the RM task.

**Figure 3.**
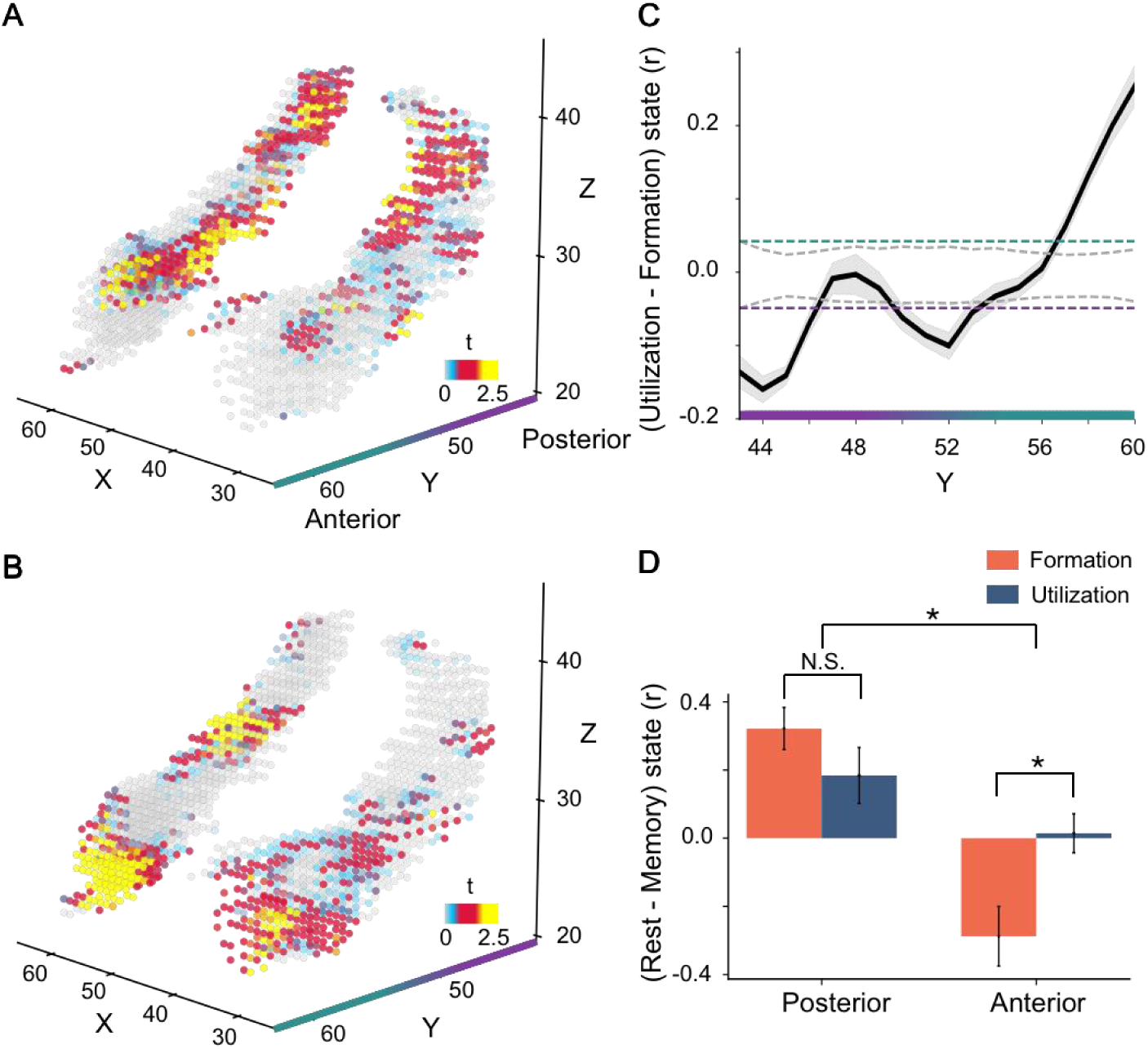
Connectivity gradient in the HPC. (A) Difference in FC of the formation state relative to the utilization state. Pearson’s correlation was performed between the mean BOLD signal series within the autobiographic network and that in each voxel of the HPC for the formation and utilization states, respectively. The difference in FC was indexed by t-value. (B) Difference in FC of the utilization state relative to formation state. (C) Difference in FC of the utilization state relative to the formation state along the hippocampal long axis. (D) FC strength of memory state (formation versus utilization) and hippocampal gradient (posterior versus anterior) compared to the rest state revealed a significant interaction. *: *p*< 0.05; Error bars: standard error.

### The causal link between functional and anatomical gradients

The findings from both representational similarity and functional connectivity analyses suggest that the posterior HPC is predominantly involved in memory formation and the anterior HPC in memory utilization. Interestingly, these function and connectivity gradients coincide with the anatomical gradient of the HPC, where the posterior HPC is characterized by a higher density of place cells with smaller receptive fields, and the anterior HPC by place cells with larger receptive fields. Specifically, previous studies have shown that the posterior HPC is preferentially associated to both short- and long-term learning (Fernandez et al., 1998; Giovanello et al., 2009; Woolley et al., 2015; Zeithamova & Bowman, 2020; Maguire et al., 2000; Weisberg et al., 2019; Weisberg & Ekstrom, 2021), implying its role in encoding information into an abstract, fine-grained representation. In contrast, the anterior HPC is often co-activated with visual sensory regions during autobiographical memory retrieval (Cabeza & St Jacques, 2007; McCormick et al., 2015; Mazzoni et al., 2019), implying its involvement in computational processes of visual action at a coarse precision. Therefore, if there is a causal anatomical-functional link, we would expect the memory formation to require a fine-grained precision, and the memory utilization to favor a coarser precision.

To test this hypothesis, we developed a hybrid Convolutional Long Short-Term Memory (C-LSTM) neural network to simulate how information is encoded and then retrieved during the RM task (Fig. 4A). The C-LSTM comprises three components: (1) a convolutional layer for encoding the visual presentation of *Good* and *Bad* cue sequences, (2) an LSTM layer for abstractly storing the spatiotemporal locations of these cues, and (3) a transposed convolution layer for predicting the location of the next *Good* cue through *Self* marker placement. A series of the C-LSTMs, varying the kernel sizes of the convolutional layers (representing visual-action resolution, VR) and the hidden sizes of the LSTM layer (representing memory resolution, MR), were trained. In this context, the MR parameter corresponds to the resolution necessary for memory formation, and the VR parameter to the resolution required by memory utilization. Our aim was to determine which combinations of VR and MR yield optimal performance of the C-LSTM in the RM task.

**Figure 4.**
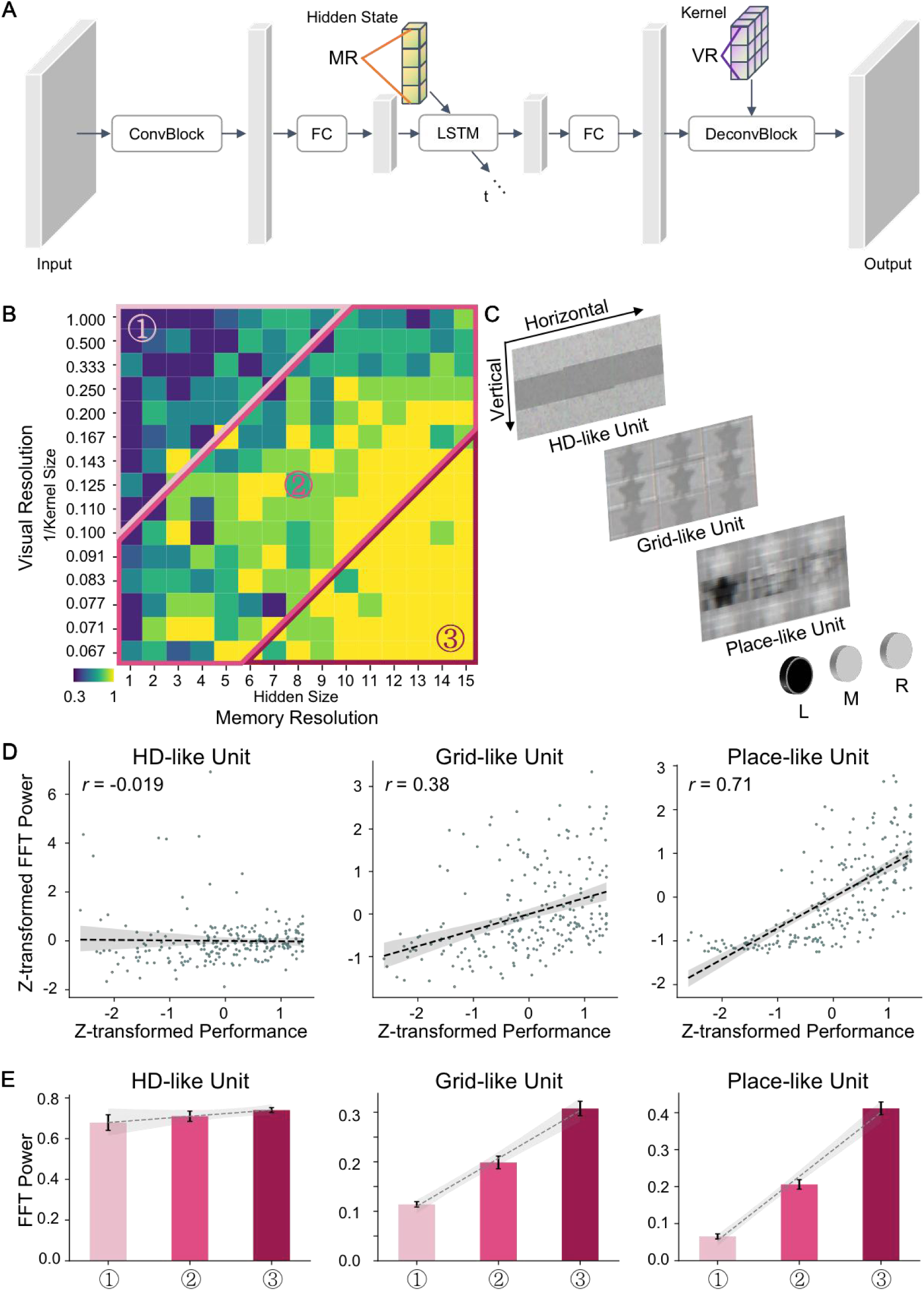
Computational modeling of memory formation and utilization. (A) Architecture of the hybrid neural network (C-LSTM). The C-LSTM comprises convolutional layers with an LSTM. The input to the C-LSTM is the visual stimuli from the RM task, and the output is an image predicting the location of the next *Good* cue (denoted by a star). ConvBlock, FC, and DconvBlock denotes convolution layer, fully connected layer, and deconvolutional layer, respectively. MR denotes the size of the hidden state in the LSTM, while VR denotes the kernel size of the deconvolutional layers. (B) Behavioral Performance of C-LSTM. The performance varied significantly across combinations of MR and VR. Each cell in the matrix indicates model performance for a specific pair of VR and MR, color-coded for prediction accuracy (dark blue: poor; light yellow: optimal). Models outlined in color were chosen for further analyses in (E). (C) Exemplar feature maps from deconvolutional layer units. The feature maps of these units were analogous to the activation patterns of biological head direction cells (top), grid cells (middle), and place cells (bottom). (D) Pearson correlations between FFT power and model performance across unit types. (E) Averaged FFT power across unit types. The presence of the HD-like, grid-like, and place-like units in models with poor ①, medium ②, and optimal ③ performance. Dashed black line denotes regression line. Shaded gray area denotes SE. Error bars: standard error.

Computational simulations revealed that both the MR and VR parameters significantly modulated C-LSTM’s performance, which was characterized by the prediction accuracy of *Good* cue location. Specifically, the performance increased with the increase of the MR parameter (*F*(14, 195) = 13.56, *p* < 0.001) and as the decrease of the VR parameter (*F*(14, 195) = 4.39, *p* < 0.001). Additionally, the interaction between MR and VR was also significant (*F*(1, 195) = 1.31, *p* < 0.05), suggesting that the increase of MR elevated the slope of the influence of VR on performance. Therefore, optimal performance was reached by a C-LSTM with a high MR value combined with a low VR value (Fig. 4B), aligning with our prediction that memory formation requires a fine-grained representation, while memory utilization prefers a coarse representation.

To further unveil how spatiotemporal information was encoded and then retrieved in our model, we extracted the feature maps of each single unit in the transposed convolution layer, because their responses directly determine model’s decision. For each unit, its weight was retained while the weights of all other units were set to zero. Accordingly, the output of the deconvolutional layer yielded a feature map of this single unit. Interestingly, the feature maps of deconvolutional units embedded the spatial structure of visual stimuli in either the vertical dimension (Fig. 4C, top), the horizontal dimension (Fig. 4C, bottom), or both the vertical and horizontal dimensions (Fig. 4C, middle). These hierarchical activity patterns of neural units are analogous to the place cells, head direction (HD) cells, and grid cells as observed in biological brains (O’Keefe & Conway, 1978; Taube et al., 1990; Hafting et al., 2005). Specifically, place-like units showed selective activation for the horizontal target locations, HD-like units responded with a band-like pattern across the rows vertically across the space, and grid-like units exhibited the square-like activation patterns tessellating the entire space (Fig. 4C). To quantitatively identify these three types of units in the transposed convolution layer and their roles on model decision, we first applied Fast Fourier Transform (FFT) to identify the spatial frequency of the feature map of each unit from 225 C-LSTMs (i.e., all possible pairs of 15 VRs with 15 MRs). Specifically, grid-like units are characterized by the minimum spectral power at 3-cycle from horizontal and vertical axes, HD-like units was expected to show the spectral power at 1-cycle in the vertical axis, and place-like units would exhibit the spectral power at 1-cycle in horizontal axes. Second, we equally partitioned C-LSTMs into three group (i.e., ①②③) according to their performance varied along the diagonal of VR-MR matrix (Fig. 4B). In this way, the participation of the three types of units can be accessed by the group of models from poor to optimal performance. As the result, significant positive correlations were revealed for the number of either grid-like or place-like units with model performance (Fig. 4D; grid-like unit: *r*= 0.38, *p*< 0.001; place-like unit: *r* = 0.71, *p*< 0.001). Correspondingly, the FFT power of these two types of unit significantly increased with model performance (Fig. 4E; grid-like unit: *F*(2, 132) = 70.02, *p*< 0.001; place-like unit: *F*(2, 132) = 177.72, *p*< 0.001). However, no significant variation of the FFT power was found for HD-like units among the models with diverse range of performance (Fig. 4D & E; *r*= -0.019, *p>*0.05; ANOVA: *F*(2, 132) = 1.26, *p*= 0.29). Therefore, the HD-like units appeared consistently across models, regardless of performance (Fig. 4E, left); however, the presence of grid- and place-like units decreased in models with medium performance and was absent in those with poor performance (Fig. 4E, middle & right). This pattern suggests a hierarchical emergence in relational memory processing, with HD-like units appearing first, followed by grid- and place-like units (see also [Wills & Cacucci, 2014] for similar patterns in mice during the development).

Taken together, our computational model demonstrates a causal link between the multi-scale representations in the HPC and the cognitive states of relational memory. Importantly, the emergence of grid-like, HD-like, and place-like units in the C-LSTM implies the essential role of the HPC in processing relational memory, aligning with observed biological phenomena (O’Keefe & Conway, 1978; Taube et al., 1990; Hafting et al., 2005).

## Discussion

The hippocampus acts as a bridge between our past and future, enabling us to learn from previous experiences and utilize that knowledge to navigate future challenges (Schacter & Addis, 2007; Addis et al., 2007; Brown et al., 2016; Gershman, 2017; Josselyn & Tonegawa, 2020; Namboodiri & Stuber, 2021). In this study, we used fMRI to investigate how the hippocampal multi-scale representation bridges the two key states of relational memory: formation and utilization. Our study revealed a functional gradient along the anterior-posterior axis of the HPC, characterized by representational similarity and functional connectivity with the autobiographical network. Specifically, the posterior HPC was predominantly involved in memory formation, while the anterior HPC was more actively engaged in memory utilization. Computational modeling of relational memory further established a causal link between the functional gradient and the HPC’s well-documented anatomical gradient, as evidenced by the size of place fields progressively decreasing along the anterior-posterior axis (Kjelstrup et al., 2008; Brun et al., 2008; Brunec et al., 2018). That is, optimal performance in relational memory tasks likely derived from the fine-grained representation of past experiences by the posterior HPC and the coarser representation of abstract knowledge for future planning by the anterior HPC. Taken together, our study not only discovered the HPC’s functional gradient for representing relational memory but also demonstrated its origin from the anatomical gradient of place cells, hereby echoing a fundamental principle in neuroscience that structure determines functionality (Suárez et al., 2020; Bassett & Sporns, 2017; Babaeeghazvini et al., 2021).

The functional division of labor in the HPC has been extensively studied, evidenced by the involvement of partial hippocampal volumes in a specific memory task. For example, the retrieval of autobiographical memory selectively activated the anterior HPC (Cabeza & St Jacques, 2007; Giovanello et al., 2009; Huijbers et al., 2011; Raichle, 2015; McCormick et al., 2015; Mazzoni et al., 2019; Smallwood et al., 2021), whereas in long-term learning, only the posterior HPC showed functional and structural alterations in response to the encoding of spatial knowledge (Maguire et al., 1997; Maguire et al., 2000; Maguire et al., 2006; Weisberg et al., 2019; Weisberg & Ekstrom, 2021). Our study further extended these findings by incorporating both memory formation and utilization in one experimental paradigm, where human participants learned spatiotemporal sequences (i.e., memory formation) and then utilized the learned relations for decision making (i.e., memory utilization). This paradigm is proved efficient to study these two states of relational memory, as evidenced by the behavioral performance displaying a piecewise-like pattern with a turning point nicely separating these two states. Further, with two neural markers of representational similarity within the HPC and functional connectivity between the HPC and the autobiographical network, a functional gradient of the HPC was discovered, with the posterior HPC primarily involved in memory formation and the anterior HPC preferentially engaged in memory utilization.

Our computational modeling further linked this functional gradient to the anatomical gradient, evidenced by physiological recordings showing a decrease in the size of place fields along the anterior-posterior axis of the HPC (Poppenk et al., 2013; Collin et al., 2015; Robinson et al., 2015; Brunec et al., 2018). By directly manipulating the size of the hidden state in the LSTM (i.e., the resolution for memory formation) and the kernel size of deconvolutional layers (i.e., the resolution for memory utilization), we explored the parameter space to identify the model that yielded the best performance. This blind exploration led us to a model exhibiting optimal performance in the relational memory task, characterized by a large scale of deconvolutional kernel size (i.e., low VR) and a small scale of LSTM hidden size (i.e., high MR). This observation provides a mechanistic insight into the alignment of the functional and anatomical gradients. That is, during memory formation, smaller sizes of place fields may facilitate the construction of high-precision abstract knowledge. In contrast, during memory utilization, larger receptive fields may facilitate the prioritization of action-related features over irrelevant ones, possibly through a mechanism of biased competition (Desimone & Duncan, 1995; Reynolds et al., 1999). Therefore, the functional and anatomical gradients in the HPC apparently reflect a computational principle tied to the multi-scale representation by place cells (Poppenk et al., 2013; Collin et al., 2015; Robinson et al., 2015; Brunec et al., 2018).

Hippocampal place cells are renowned for their pivotal role in creating a cognitive map of an environment by encoding specific locations and directions during navigation (Schiller et al., 2015; Epstein et al., 2017). Our study extends these known functionalities, proposing their significant contribution to both the formation and utilization of relational memory. The integral role of place cells aligns with the concept that the HPC provides a unified framework where spatial memory and episodic memory rely on relational representations that integrate spatial localization with temporal continuity (Cohen & Eichenbaum, 1993; Eichenbaum, 2000; Eichenbaum & Cohen, 2014). Consistent with this speculation, our analysis of the feature maps of single units in the deconvolution layers revealed the presence of place-like units, as well as HD-like and grid-like units. Critically, the absence of these units, especially the place-like and grid-like units, significantly impaired the model’s performance in the relational memory task. These self-emerged units in our computational model, analogous to their biological counterparts essential in navigation, may constitute the implementational basis for relational representations. Therefore, our findings not only shed light on a broader application of hippocampal place cells in relational memory processing but also reinforce the concept of a unified framework underpinned by these specialized cells in the hippocampus (Epstein et al., 2017; Whittington et al., 2022; Sorscher et al., 2023).

There are several limitations warranting further investigation. First, our study suggests a continuous functional gradient along the anterior-posterior axis of the HPC, contrasting with the view of distinct functionalities for the posterior and anterior regions. However, this functional gradient, although in alignment with the anatomical gradient in the size of place fields, may also be influenced by our experimental paradigm. In the RM task, memory formation and utilization were intertwined; for example, elements of memory utilization were present even during the formation state due to task requirements. Similarly, in our fMRI data analysis, the BOLD signal series were bifurcated into two equal-length segments, the first half as the formation state and the second half as the utilization state, primarily because a distinct turning point at the trial level was not prominent. This intermingling nature of the formation and utilization states may contribute to the observed gradient in both representational similarity and functional connectivity. Future research is needed to more distinctly decouple these two states to better depict the functional gradient of the HPC. Second, the memory formation and utilization states in our study are markedly different from the states of retrieving the past and imagining the future, which are associated with scene construction (Gaesser et al., 2013) and engage both the anterior and posterior HPC (Strange et al., 2014). Future research should aim to characterize relational memory from multiple dimensions, such as spatial and temporal dimensions, and then integrate these dimensions to comprehensively understand relational memory in the hippocampus, hereby offering deeper insights into the multifaceted functional roles of the HPC. Third, while our model successfully simulated the hippocampal multi-scale coding at behavioral level, it is important to recognize its limitations in terms of biological plausibility, especially concerning its anatomical implementation. Future research should aim to develop models that more accurately resemble the HPC’s actual anatomical structure, thereby bridging the gap between computational modeling and biological mechanisms of the HPC.

## Acknowledgments

The work was supported by China Postdoctoral Science Foundation (2022M710470, B.Z.), Beijing Municipal Science & Technology Commission & Administrative Commission of Zhongguancun Science Park (Z221100002722012), and Double First-Class initiative Funds for Discipline Construction of Tsinghua University (J.L.).

## Author contributions

J.L. conceived the study. Y.L. developed the task. W.D and Y.L collected the behavioral and fMRI data. W.D. performed the data analyses under B.Z.’s supervision. B.Z. developed the C-LSTM network. B.Z. and W.D. performed the network simulation. W.D., B.Z. and J.L. wrote the paper.

## Data availability

The data that support the findings of the present study are available from the corresponding author (J.L.) upon request.

## Code availability

All analyses reported in the present study were conducted with the customized codes written in Python (version 3.9) and Matlab (version 2021a). The codes will be available from the corresponding author (J.L.) upon request. The neuroimaging toolbox and machine learning framework used by the present work are publicly available as following: FSL (version 6.0, https://fsl.fmrib.ox.ac.uk/fsl/fslwiki), BrainNet Viewer (version 1.7, https://www.nitrc.org/projects/bnv/), and Torch (version 1.9, https://pytorch.org/).

## Methods

### Participants

Sixteen participants (mean age 23.00 ± 2.97, 9 female) with normal or corrected-to-normal vision were recruited from Tsinghua University. Informed consent was obtained from each participant before participation in accordance with the experimental protocol approved by the Institutional Review Board of Tsinghua University.

### Relational Memory (RM) task

The experimental paradigm was a modified version of a sequential decision making task (Insa-Cabrera et al., 2011; Tartaglia et al., 2017). In each trial, a *Good* and a *Bad* cue were pseudo-ramdomly presented on two out of three horizontal positions of screen after a 10-s fixation. The position of the cues independently varies along 30 consecutive steps. The *Good* and *Bad* cue were never placed in the same position in sequence. Participants were required to predict the next position of the *Good* cue by placing a Selfmarker “avatar” at the anticipated location, and avoid the position of *Bad* cue during 1700 ms. After a self-paced response, feedback is immediately presented for 300 ms with the positive feedback indicated by a green upward arrow and negative feedback represented by a red downward arrow. If the chosen position is neither *Good* nor *Bad*, neutral feedback will be presented. Participants received +1, -1, and 0 virtual coin after positive, negative, and neutral feedback, respectively. If no response was made, the next step will be presented after 2000 ms. Participant’s response, feedback, and reaction times were reported during experiment.

The experiment was conducted in two consecutive days. Participants performed the task in behavioral room on day 1 while they completed the task under MRI scanning on day 2, respectively.

The behavioral task employed a four-position design programmed by PsychoPy (version = 2022.1.0), where the length of sequences (i.e., numeric strings) varied from 2 to 8 steps with a step size of 1 and eight repetitions. This design resulted in 56 trials. The total duration of experiment took approximately one hour. Experimental stimuli were presented by a 15.4-inch MacBook Pro with a screen resolution of 2880 x 1800 pixels. Participants completed six practice trials prior to the experiment to ensure that they fully understood the instructions.

The fMRI task was designed with three positions. The length of numeric strings varied from 2 to 4 steps with a step size of 1 and eight repetitions, resulting in a total of 24 trials, which were randomly assigned to 4 scanning sessions with 6 trials each session. The total duration of fMRI experiment took approximately one hour. The experimental stimuli were presented using a visual/audio stimulation system (Shenzhen Sinorad SA-9939) with a customized-design (Shenzhen Sinorad Medical Electronics Co., Ltd) MRI compatible 40” LED liquid crystal display. The distance between participant and the display was 100 centimeters. The screen resolution was 1024 x 768 pixels. Participants viewed the stimuli with an angled mirror mounted on the head coil.

### MRI Acquisition

BOLD MRI images were acquired using a 3T Siemens Prisma scanner equipped with a 64-channel phased-array head coil. Functional data were acquired with a gradient-echo-planar T2* sequence (TR: 2000ms; TE: 34.0ms; matrix size: 100×100×72; flip angle: 90°; resolution: 2 × 2 × 2 mm^3^; FoV = 200mm; number of slices: 72; slice thickness: 2 mm; slice orientation: transversal). T1-weighted three-dimensional anatomical images were collected to facilitate registration (MPRAGE: TR: 2530 ms; TE: 2.27 ms; matrix size: 208×256×256; flip angle: 7°; resolution: 1×1×1mm^3^; number of slices: 208; slice thickness: 1 mm; slice orientation: sagittal).

### fMRI Pre-processing

T1-weighted structural and functional images were analyzed using FSL (FMRIB’s Software Library, version 6.0, http://fsl.fmrib.ox.ac.uk/fsl/; Woolrich et al., 2009; Smith et al., 2004; Jenkinson et al., 2012). The non-brain tissues of high-resolution T1 images were removed using BET (Smith, 2002). The BOLD signal series of each scanning session was preprocessed independently using FEAT (FSL’s FMRI Expert Analysis Tool, version 6.0; Woolrich et al., 2001; Woolrich et al., 2004), where motion correction, slice-timing correction, and high-pass filtering at 100 s were performed. For representational similarity analysis (RSA), BOLD signal series was left unsmoothed to preserve fine-grained spatial information (Chadwick et al., 2012). For functional connectivity (FC) analysis, BOLD signal series were smoothed using a Gaussian kernel of full width at 5mm half-maximum (FWHM). These preprocessed functional images of each participant were normalized to standard Montreal Neurological Institute (MNI-152) template using FSL FLIRT (Jenkinson & Smith, 2001; Jenkinson et al., 2002) before being sent to group-level statistics.

### Representational similarity analysis

The functional separation of the HPC was examined by the representational similarity analysis (Kriegeskorte et al., 2006; Kriegeskorte et al., 2008). First, the BOLD signal series of task period were subdivided into two temporal intervals with equal duration, which were labelled as the relational memory formation state and the utilization state, respectively. Second, a general linear model (GLM) was created for fitting the BOLD time series of each voxel, where 18 regressors were specified. These regressors indicated the cognitive states corresponded by the rest, formation, and utilization periods from each of the six trials, and the six motion parameters were additionally included as covariates. Third, the multivoxel patterns were extracted from the beta images using a 6mm radius sphere centered at each voxel of the HPC. Pair-wise Pearson correlations were performed across the 18 cognitive states. This procedure resulted in a 18 × 18 correlation matrix, which was Fisher’s r-to-z transformed. Fourth, two additional GLMs were created to examine the neural representation of memory formation and utilization states, respectively. Specifically, a binary regressor was used for each GLM to fit the correlation coefficients of cognitive states, where 1 and 0 denoted “same” and “difference”, respectively. A higher beta coefficient of the two GLMs in any voxel indicated stronger representational similarity for the memory formation and utilization states. Then, the representational similarity maps were transformed into MNI space, averaged across sessions, and finally sent to second level analysis. Identical procedure was applied to the RSA of the rest and task periods.

### Functional connectivity analysis

To examine the connectivity separation of the HPC, we computed the connectivity pattern of the HPC for each of the memory formation and memory utilization states. Given the brain areas within the autobiographical network intrinsically connect to the HPC (Conway et al., 2001; Spreng & Grady, 2010; Sestieri et al., 2011; Philippi et al., 2015; Smallwood et al., 2021), we particularly focused on the connections between the autobiographical network and the HPC. First, we derived the functional mask of the autobiographical network by performing a contrast of the hippocampal whole-brain connectivity map from the task period to the rest period (Fig. S2). Specifically, a GLM was created to regress out the potential nuisance factors, which included the six parameters derived from rigid body correction of head motion (Birn et al., 2006; Wise et al., 2004), the mean time courses within the lateral ventricles, and the mean time courses within the white matter (Bartels & Zeki, 2005; Fox et al., 2005). Second, the BOLD signal series of the task and rest periods were extracted through concatenating the TRs (i.e., 5 TRs) across trials (i.e., 6 trials) independently for each session. Third, the whole brain connectivity map of the HPC for each of the rest and task state was computed by performing Pearson correlations between the regional time courses of the HPC (the HPC mask was obtained from the Harvard-Oxford (HO) atlas included in FSL) and the BOLD signal series of each voxel of the remaining brain. Fourth, the contrast in the connectivity of the HPC between the task period and rest period was performed after Fisher’s r-to-z transformation (Fig.S2). This result was visualized using BrainNet Viewer (version 1.42, https://www.nitrc.org/projects/bnv). The functional mask of the default mode network was then generated using the contrast map with the voxels within the medial temporal lobe excluded. Finally, the connectivity separation of the HPC was examined by computing the Pearson correlations between the regional time course of default mode network and the time course of each voxel of the HPC in each of the memory formation and utilization states, which was extracted by concatenating the TRs (i.e., 15 TRs) across trials (i.e., 6 trials). The state-dependent connectivity maps of the HPC were transformed into standard MNI152 space, averaged across sessions after Fisher’s r-to-z transformation, and sent to second level analysis.

### Convolutional LSTM (C-LSTM) model

An encoder-decoder network programmed by PyTorch (version = 1.9; https://pytorch.org/) was built to perform the RM task, where the encoder included a convolutional layer followed by a fully connected layer, which projected the flattened output to a long short-term memory (LSTM) layer. The encoder accepted the images with dimension of 90×40×3 pixels as input and constructed the memory space storing the sequential relation of experimental stimuli. The decoder included a fully connected layer followed by a transposed convolution layer, which was used to generate visual output, indicating the predicted location of the *good* cue. In other words, the decoder utilized the sequential relations stored by memory space to guide actions on visual space. To examine the role of coarse / fine-grained scales akin to the hippocampal ventral-dorsal axis on relational memory formation and utilization, we modulated the visual resolution (VR), defined by 1/kernel size of convolutional layers (ranged from 1 to 15) and the memory resolution (MR), defined by the hidden size of LSTM (ranged from 1 to 15) to examine their influence on model performance. The kernel size of the convolutional layer was set the same as that of the deconvolutional layer based on the biological basis of the visual cortex (Miyawaki et al., 2008). In practice, the kernel size could vary independently, and the variance of the kernel size of the convolution layer had little effect on the findings (Fig. S3). During training, the Adam optimizer with a learning rate of 0.001 and a weight decay parameter of 1e-5 were used. The optimization process operates within a batch size of 8, which equals to the length of sequential images and spanned 1,000 epochs for each kernel size of convolutional layer and the hidden size of LSTM. The loss function is specified as Mean Squared Error Loss. To improve the efficiency of network training, the encoder was independently trained in advance with a fully connected layer as decoder in predicting the upcoming location, then C-LSTM used the pretrained weights of encoder, which was fixed during training.

### Statistics

The behavioral analysis was performed using Python 3.9 and SPSS 20. The second level analysis of RSA and functional connectivity analysis were performed using two tailed one-sample t-test. The generated t-maps used an initial threshold of *p* = 0.01. If no clusters were revealed, a liberal threshold of *p* < 0.05 was used. The reliability of the uncorrected clusters was further examined using a non-parametric statistical inference that does not make assumptions about the distribution of the data (Nichols & Holmes, 2002). Specifically, 5,000 random sign-flips were performed on the parametric images of RSA and functional connectivity. The clusters with size higher than 95% of the maximal suprathreshold clusters from permutation distribution were reported. Same permutation procedure was applied to evaluate the statistical model fitness of participant behavior, the variation of the representational similarity and functional connectivity along hippocampal long axis, and the identification of unit type in C-LSTM. The main effect of C-LSTM parameter VR and MR and their interaction on task performance was examined by ANOVA. The unit type of C-LSTM was determined by computing the spatial frequency in both the horizontal and vertical dimension from the feature maps using Fast Fourier Transform (FFT). The linear relationship between the unit type and model performance was examined by Pearson correlation. ANOVA was employed to examine the significance of the FFT power variation of unit types relative to model performance.

## Supplementary

**Figure S1.**
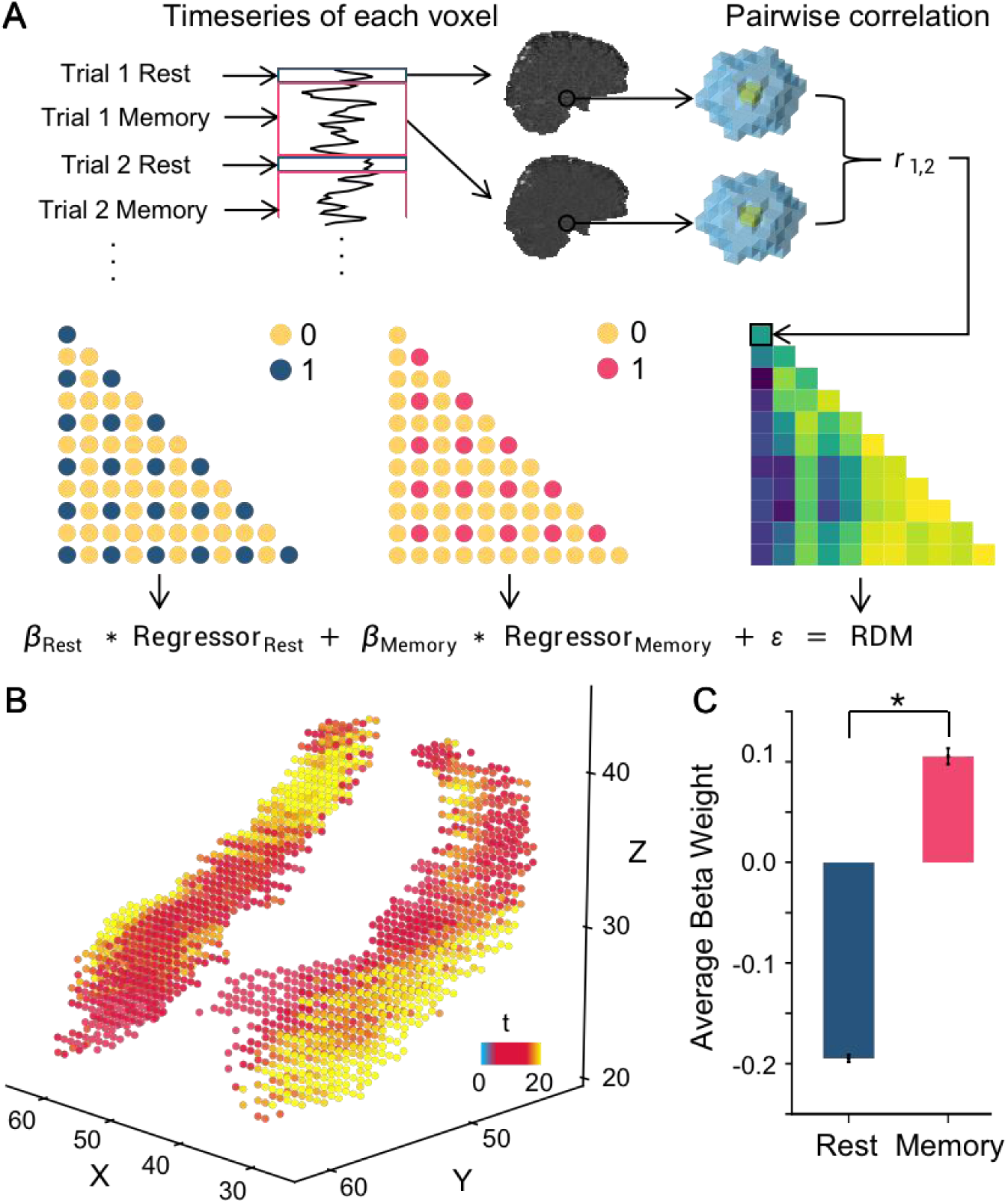
Representational similarity of rest and memory state. (A) The schema of RSA analysis. First, the multi-voxel patterns of the rest and memory state were derived by a GLM. Second, the 12×12 similarity matrix, built through the searchlight-based Pearson correlation of multi-voxel patterns among trials, was fitted by another GLM, which specified the cognitive state using binary regressors. The blue and red points in the binary matrix denote “same” between trial-pair. The estimated parameters of GLM were used for the analysis of (B). (B) The neural representation of memory relative to rest state. The value in each voxel represents t-value. The Y-axis corresponds to the coordinates of hippocampal long axis. (C) ROI analysis. The HPC showed significantly higher similarity in memory state relative to rest state (*t*(15) = 33.45; *p*< 0.001, two-tailed).* denotes the threshold of *p*< 0.05. Error bars denote standard error.

**Figure S2.**
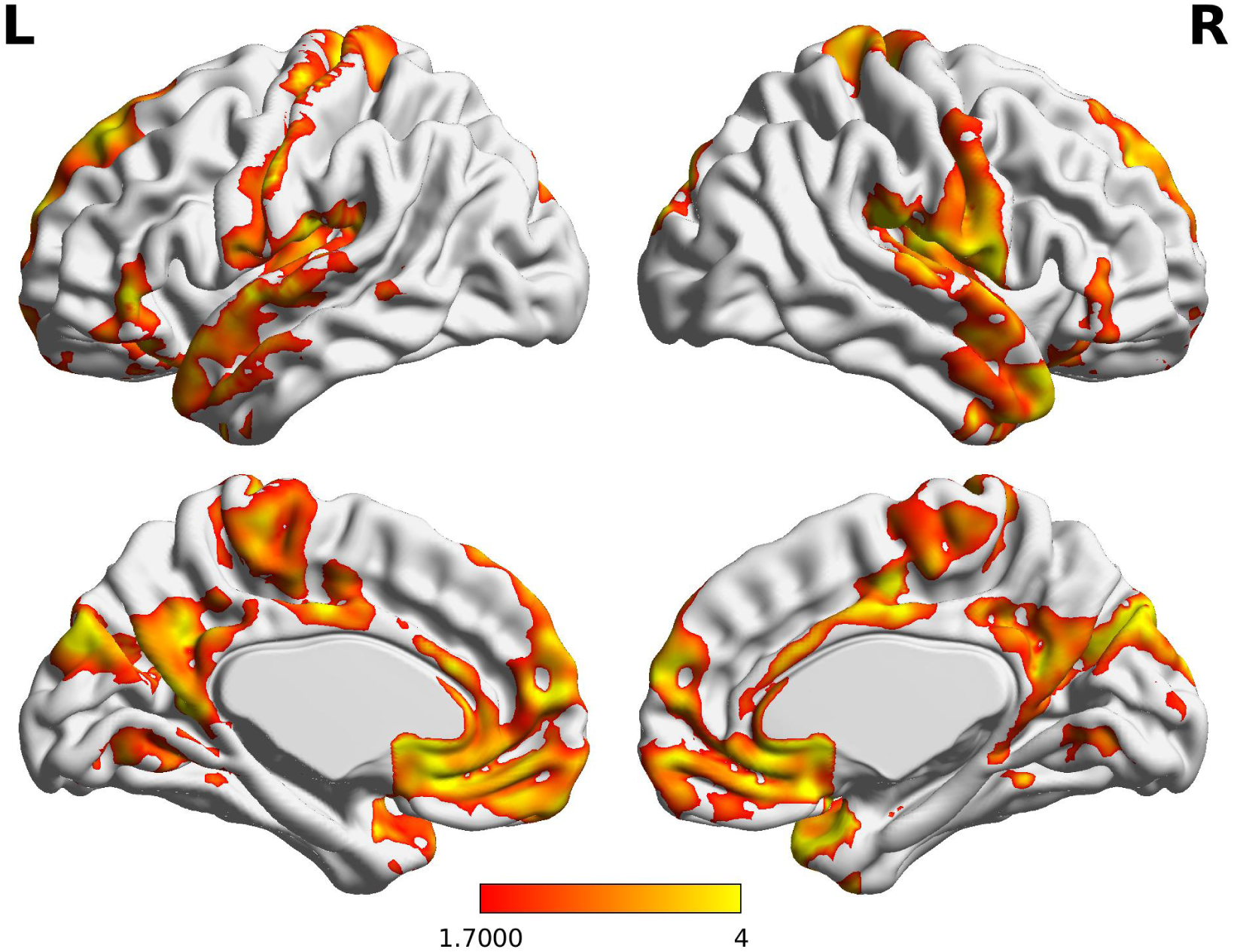
Functional connectivity of HPC in memory state compared to rest state. The functional connectivity was computed for each participant in both the memory and rest states. A Pearson correlation was performed between the mean BOLD signal series within the HPC and the voxel in the rest of brain. A contrast analysis was conducted comparing memory state relative to rest state. All clusters were corrected by False Discovery Rate (FDR) with a significance threshold of *p*< 0.05. colorbar denotes *t* value.

**Figure S3.**
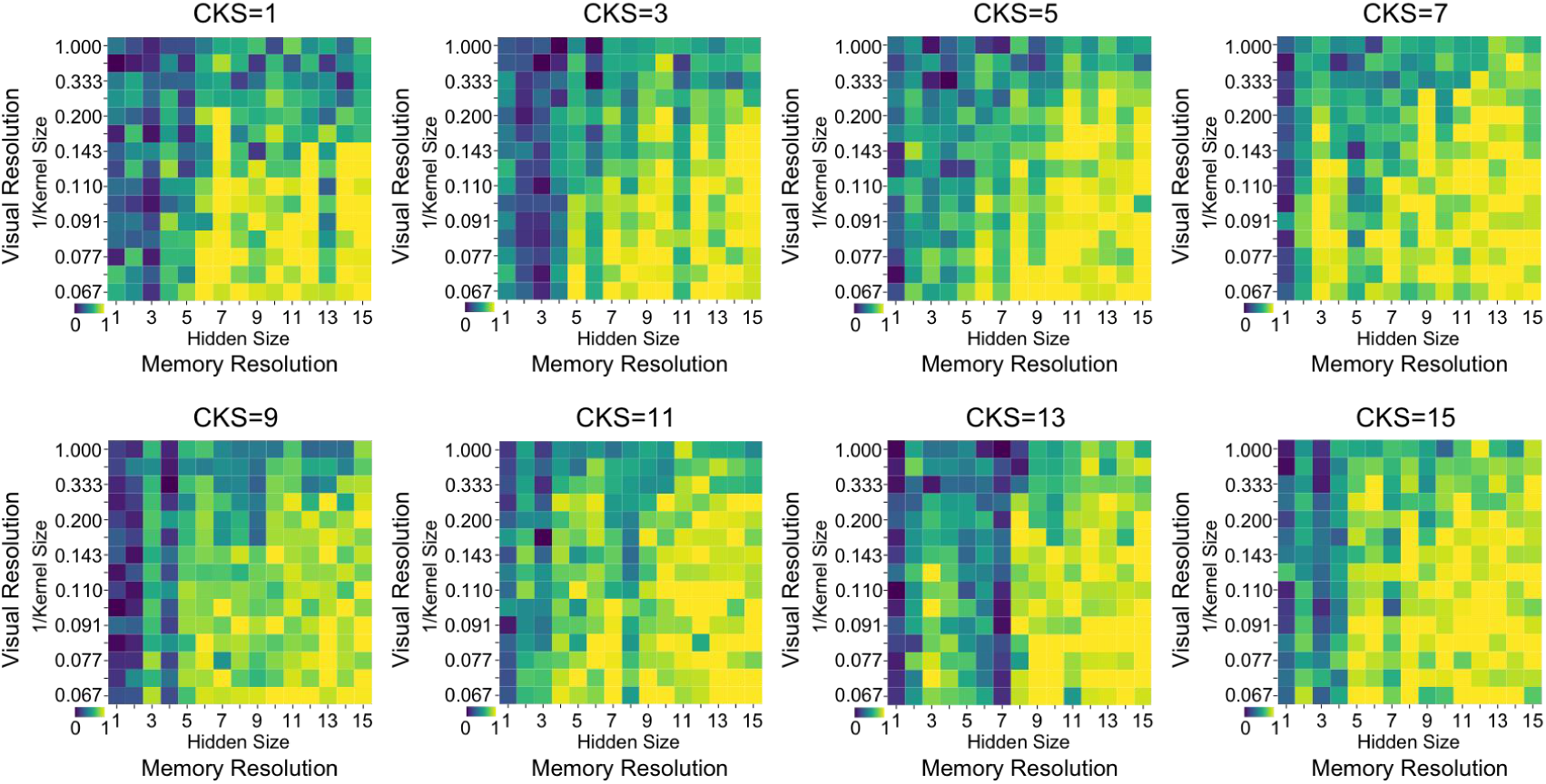
Behavioral performance of C-LSTM with fixed convolutional kernel size (CKS). Across a wide range of CKS values (i.e., varying from 1 to 15 in steps of 2), consistent patterns similar to Fig. 4B emerged. That is, optimal performance was achieved with smaller VR values (i.e., coarse representation during memory utilization) and larger MR values (i.e., fine-grained representation during memory formation). The color coding follows the scheme in Fig. 4B.

